# Global brain circuitry control of behavior emerging from self-governing vector field dynamics in subnetworks

**DOI:** 10.1101/2025.07.30.667657

**Authors:** Szilvia Szeier, Henrik Jörntell

## Abstract

Behavior ultimately depends on the spatiotemporal patterns of the neuron population activity across the brain. Here we address the issue of how the evolving patterns of population activity can be governed by the intrinsically available mechanisms within the brain. We show how the control of the evolving activity can be represented by a high-dimensional vector field, which is an emergent effect of the integrative effects afforded by the membrane capacitance of the neurons and the weights of the synaptic connections between them. For each subnetwork of the brain, its intrinsic connectivity defines the structure of a lower-dimensional vector field with a local attractor point, towards which the subnetwork activity is constantly drawn. We show that other subnetworks, defined by having a degree of independence from but an impact on the first subnetwork, will constantly move the location of the local attractor point, causing the activity in the first subnetwork to constantly ‘chase its own tail’. We show how this principle can explain how minor differences in the corticospinal control signal can produce a variety of movement patterns through the spinal interneuron circuitry. At the global level, we show how the cortical neuron population can be thought of as many concatenated subnetworks that produce diverse dynamic evolutions across the cortical neuron population globally. This operational principle can produce a dynamic population activity reminiscent of that observed across the brain *in vivo* and explain the foundational mechanisms underlying autonomous cortical control of its own activity evolution.

## Introduction

Technological and methodological advances have made investigations of the joint dynamic activity of large numbers of neurons much more common. A better grasp of the activity dynamics at the neuron population level has allowed us to gain a deeper understanding of brain function (Vyas et al 2020). A common analysis method for neuron population activity includes a top-down approach of training Artificial Neural Networks (ANNs), or some other selective dimensionality reduction method, to obtain a trained network structure that approximatively predicts the neuron population recording data. These lower-dimensional embeddings of the recording data have shown to be effective in specific applied domains but can also fail to generalize to out-of-domain examples (Jazayeri & Ostojic 2021). Hence, while an experimenter may apply any decoding scheme of choice to match hypothesized low-dimensional latent variables with the recording data in a certain brain area under specific task conditions, there is no *a priori* reason to assume that the brain operates like that scheme. Instead, the schemes can be parallel complex systems that are made to match the recorded activity under the given conditions (Jazayeri & Ostojic 2021, Kristensen et al 2024). But outside the constrained lab conditions, higher-dimensional aspects of the neuronal network behavior may for example be critical for the high degree of behavioral diversity, adaptability and flexibility observed in biological systems (Kristensen et al 2024, Musall et al 2019, Yin et al 2025). The only exception when approximative models of brain activity would work in a more general sense would be for the one true dimensionality reduction method which reflects how the biological brain operates at the circuitry level, which has yet to be discovered. Therefore, our aim here was not to approximate recording data but instead to investigate the neural mechanisms underlying the generation of neuron population dynamics, focusing on the emergent effects of the brain’s intrinsic network operation in its most generic form.

The spatiotemporal pattern of the neuron population activity directly underlies behavior, since behavior ultimately consists in spatiotemporal patterns of muscle activity driven by these neuron populations. This behavior can manifest itself in multiple forms, for example in overt forms of behavior, which is observable from the outside, but also in internal, non-observable, state changes in the network. These would correspond to what we might otherwise refer to as “thoughts”, where the brain integrates sensory information across multiple time scales to control future behavioral decision making. Here we build on our previous work of how the network connectivity results in a vector field structure that controls the activity distribution across the neuron population (Szeier & Jörntell 2025) and extend the analysis into the temporal dynamics domain. We begin by demonstrating that the combination of membrane capacitance and constituent static leak channels in the neurons gives rise to attractor points in the vector field, towards which the network activity inevitably converges, regardless of its initial activity state. To achieve dynamic activity evolution resembling that observed in the brain, we show that the network must incorporate additional partially independent neural components so that one subnetwork controls the attractor point and the vector field of another subnetwork. This is shown to result in a continually evolving attractor point in the targeted subnetwork and thereby a constantly evolving activity distribution across its neuron population, reminiscent of observations in the brain *in vivo* (Etemadi et al 2022, Norrlid et al 2021, Stringer et al 2019). We show in principle how this mechanism can account for corticospinal control of the vector field in the spinal cord circuitry, enabling the generation of various dynamically altering spatiotemporal patterns of muscle activity, and how the same principle can be applied to understand the fundamental mechanism emerging into the intrinsic self-governing of cortical activity dynamics.

## Methods

### Conductance-based non-spiking neuron model

To investigate neuron population dynamics, we built network models using a conductance-based, non-spiking neuron model first introduced in (Rongala et al 2021). This model, denoted the Linear Summation Neuron Model (LSM), encompasses a linear summation of synaptic input that then contributes to the neuron’s dynamic state transition. The activity level of the post-synaptic neuron depends on the presynaptic input(s) and can be summarized by

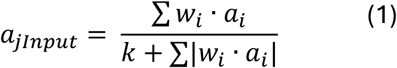

where the summation iterates over all presynaptic neurons *i* of neuron *j*. The activity level of presynaptic neuron *i* is indicated by *a_i*, and the weight (or strength) of the connection from neuron *i* to neuron *j* by *w_i*. The denominator includes a factor that divides the incoming presynaptic inputs by the total currently active presynaptic activity, hence producing a shunting effect that scales with the total synaptic activity. The *k* parameter acts as a normalization which aims to reflect the constituent leak channels that give rise to the neuron’s membrane potential. Since the density of leak channel is roughly constant (Spruston & Johnston 1992, Thurbon et al 1998) and the membrane area scales with the number of synapses the neuron receives, the *k* parameter is scaled with the number of synapses.

Then, the state transition of the neuron model can be described as

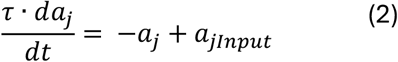

where τ is the neuron membrane time constant, which is referred to as the dynamic leak in (Rongala et al 2021), and *a_jInput* is the formula expressed in Equation 1. In a biological cell, the time constant depends on both the capacitance and the resistance (leak) of the membrane, and because of the capacitive effect the membrane time constant can be said to constitute a type of state memory in the neuron. It should be noted that in this instantiation of the LSM model, a neuron’s resting state corresponds to an activity level of zero, and the dynamic leak will over time tend to bring the neuron back towards its resting state. Whenever the activity level of the LSM neuron exceeds the resting state value, it provides synaptic input to its downstream neurons. In some cases, where explicitly defined, we instead applied a higher threshold level in subsets of the neurons.

## Results

### Temporal evolution of the network state

Network population dynamics can be encapsulated in the high-dimensional activity space where each dimension represents the activity level of a neuron in the network. Consequently, this activity space has as many dimensions as there are neurons. The location within the space uniquely defines the activity level distribution across all neurons, which corresponds to the ‘state’ of the network. Therefore, we can instead refer to the activity space as the state space of the network.

In previous work, we introduced how synaptic connections between neurons correspond to the impact exerted by the pre-synaptic neurons on the post-synaptic neuron. This impact can be thought of as a vector, which depends on the activity of the neurons in question and the strength, or weights, of their synaptic connections. For a given synaptic weight configuration, the set of all possible activity level combinations determine a vector field. The vector field geometry can be manipulated through synaptic weights, as shown in (Szeier & Jörntell 2025). This vector field impacts the direction of the temporal evolution of the network population activity, or of the activity state of the network. In an example excitatory 2-neuron network (Figure 1A), a connectivity motif which is known to exist within the cortex for example (Cossell et al 2015, Peng et al 2024), the mutual synaptic connections would tend to result in an ever-increasing activity level in the two neurons for any non-zero starting state (Figure 1B left).

**Figure 1.**
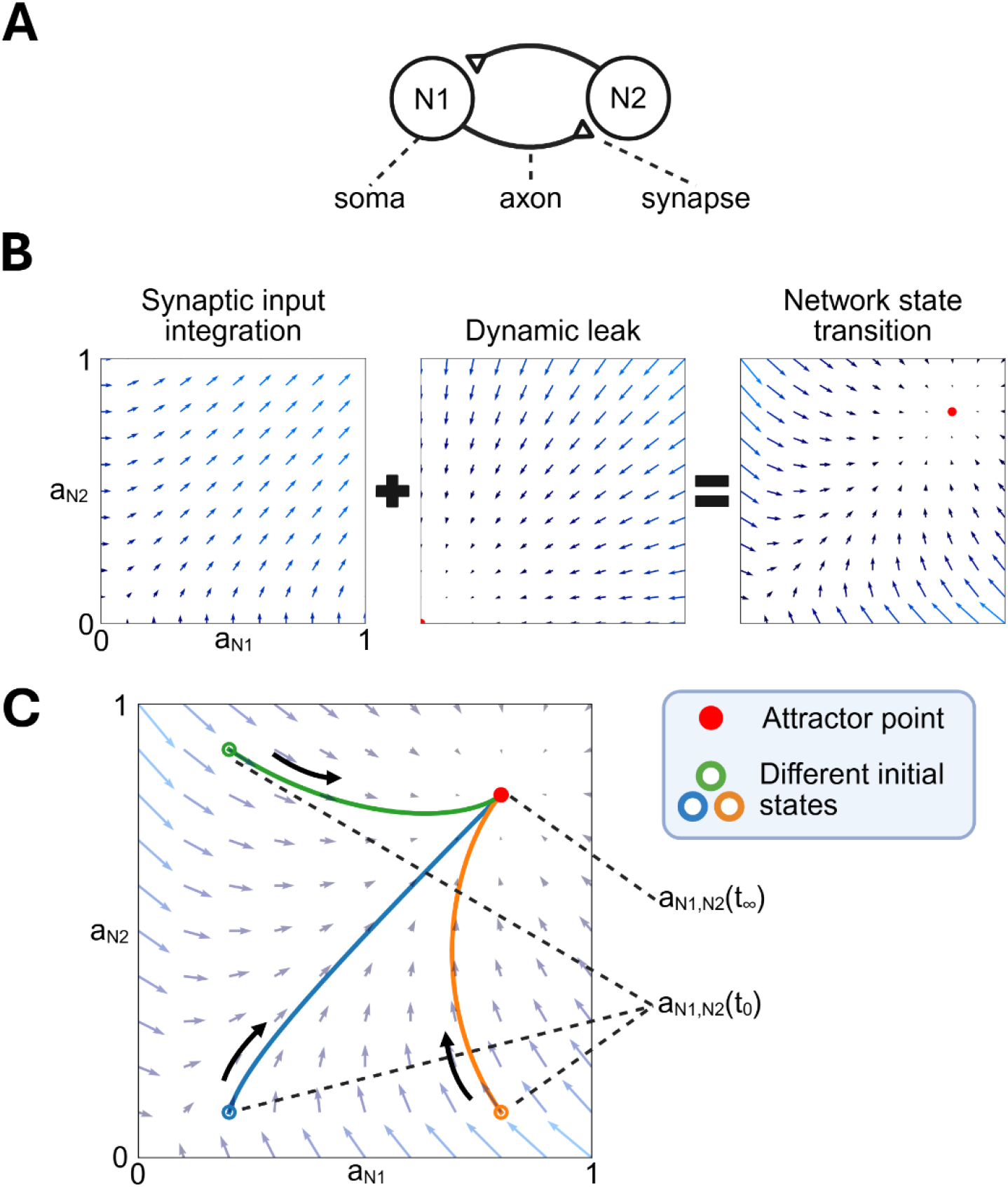
Factors impacting the temporal evolution of the network state within a fully connected excitatory neuron network. A) Network consisting of 2 excitatory neurons (white circles). The neurons impact each other through synapses, located at the axon terminals, where the synapses in this example have the same weights. B) (left) The synaptic connections constitutes a vector field (Szeier & Jörntell 2025), which drives the state of the network towards the upper extreme of the activity state space. (middle) The time constant of the membrane of the neuron (its ‘dynamic leak’ component) results in a corresponding vector field with an attractor in the origin. (right) The combination of the synaptic weights and the dynamic leak results in a vector field with a non-zero attractor point. C) The attractor point (red circle) will pull the network activity towards itself, regardless of the initial activity state. Upon reaching the attractor point, the network state will remain there indefinitely.

However, this previous work illustrated the case that applies when neurons have no state memory. If the neurons have no state memory, then the network is 100% volatile, which means that each time step is unique, a *de novo* state starting from zero. But neurons do have a state memory. The capacitor of the neuron, i.e. its membrane capacitance, contributes to a state memory, which in the case of only excitatory synaptic inputs, would be integrating consecutive synaptic inputs into a gradually increasing membrane potential level. The capacitor also impacts the rate at which a biological neuron’s activity level can change, through the resulting membrane time constant, which we in our neuron model refer to as the dynamic leak (Rongala et al 2021). The dynamic leak is an activity-dependent decay rate term that tends to bring the neuron’s activity level back to zero, thereby shunting the state memory effect imparted by the capacitor. Therefore, the dynamic leak in itself generates a vector field where all vectors point towards the origin (Figure 1B middle). Under the presence of dynamic leak, an attractor point naturally emerges at the origin (zero level) of the state space, which is in line with that a neuron’s resting membrane potential, or its resting state, acts as an attractor point for that neuron, as described already in the early Hodgkin-Huxley model of neuron function.

The contribution of the neuronal synaptic connections combined with the dynamic leak can lead to the emergence of attractor points within the neural state space that are located away from the origin (Figure 1B right). (Without the capacitor, these points would instead have been critical points, i.e. as illustrated in (Szeier & Jörntell 2025)). The combined effect of the synaptic integrator imparted by the capacitor and the dynamic leak determines the state transitions of the network (Figure 1B). In the example network (Figure 1A), starting from different initial states, the activity trajectories, or the gradual state transitions, will then take different paths but will nonetheless converge to the same point (Figure 1C), where they will remain in place perpetually if no other factor intervenes to change the attraction to that state.

In experimental recordings *in vivo*, the activity of the neuron population is, however, dynamically changing all the time (Etemadi et al 2022, Norrlid et al 2021, Stringer et al 2019). We therefore hypothesized that, in this conceptual framework, the situation where the network state can reach the attractor point needs to be avoided as it would result in stagnant activity trajectories in conflict with the *in vivo* observations. The attractor point located at the origin, which would mean that all neuron activity has stopped, should naturally also be avoided, because then the brain activity would be perpetually stopped. The changing and dynamic activity level distributions observed in brain recordings would correspond to moving activity trajectories in the neural state space, which in the present framework could only be achieved if the attractor point of the network is constantly moving. Since the attractor point is the product of the whole connectivity of the network, any movement of the attractor point could only be expected to be gradual, i.e. it would form an *attractor trajectory*.

Temporary changes in synaptic weights, such as short-term facilitation or short-term depression, could be one possibility to obtain moving attractor points, but our assumption here is that because there are only limited short-term synaptic adaptions in the brain *in vivo* (Jorntell & Ekerot 2006) we can ignore this factor as being of major importance here. More non-temporary forms of synaptic plasticity *in vivo*, i.e. LTP or LTD, will, at least over the course of minutes, only result in minimal weight changes if any at all (Jorntell & Ekerot 2002, Jorntell & Ekerot 2003, Jorntell & Ekerot 2011), and we hence consider these processes too slow to ‘rescue’ the network from getting caught in its attractor point. For these reasons we consider the weights to be static. This raises the question of what mechanisms give rise to a moving attractor point that appears to be the case in brain activity *in vivo*.

### Independence among neuron populations

To overcome the static convergence of the network in Figure 1, where the state inevitably will settle in its attractor, we used another two-neuron network to examine how external influence could disrupt or reshape the dynamics of that subnetwork. Figure 2A illustrates a subnetwork composed of an excitatory and an inhibitory neuron, which is controlled by an upstream subnetwork (without any defined structure in this example). The downstream subnetwork is here the “readout population”, i.e. the plane of the vector field, or the subspace of the whole network’s state space, that is visualized in Figure 2B-D. The activity of the upstream subnetwork is in this case fully independent of the activity in the readout population, meaning that the downstream subnetwork cannot impact the former. At each time point, the activity level in the upstream population will determine the location of the vector field plane, and thereby also the location of the attractor point that governs the dynamics of the readout population (Szeier & Jörntell 2025) (Figure 2B). Hence, from the readout population’s point of view, the attractor point will gradually move between different locations that the activity state of these two neurons will tend to follow over time (Figure 2C). The downstream population’s pursuit of the attractor point generates an activity trajectory, which will also depend on the initial state of the readout population (Figure 2D).

**Figure 2.**
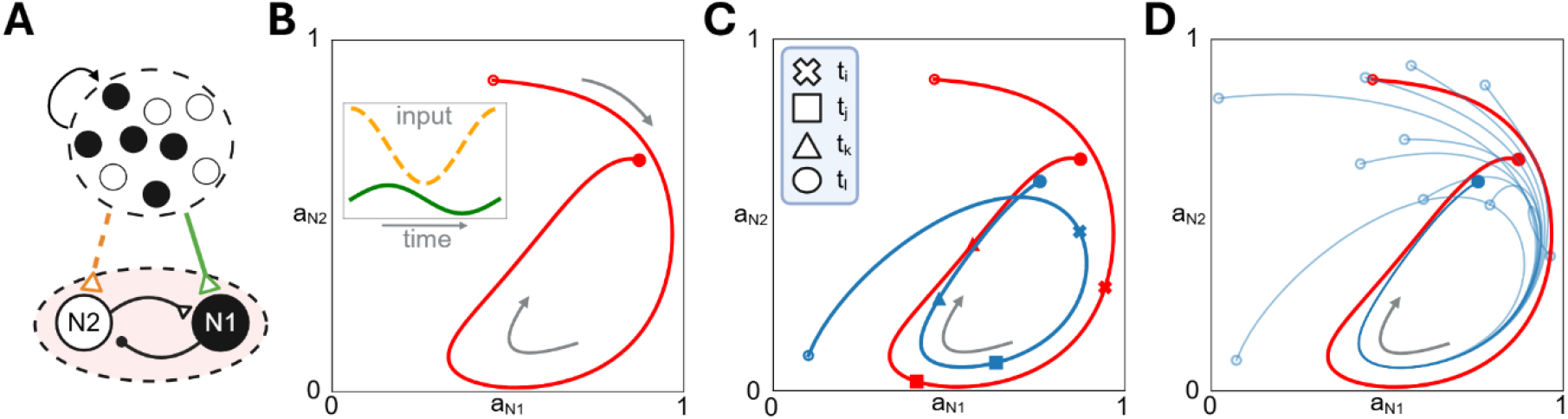
An upstream network controls the attractor point of a downstream network. A) A downstream network consisting of an excitatory and an inhibitory neuron (white and black circles, respectively) receives input from an undefined upstream network, with two outputs. B) The input received from the upstream population (inset) controls the attractor point of the downstream population, thereby creating an attractor trajectory (red curve). C) The activity state of the downstream population converges towards the moving attractor point, which results in an activity trajectory (blue curve). At different time points (symbols), the state of the downstream population tends towards but never reaches the attractor point at that point in time. D) The choice of initial state will impact the activity trajectory of the downstream population as it converges towards the attractor trajectory.

This effect emerged because the readout population lacked synaptic connections to the upstream subnetwork, whereas the upstream subnetwork influenced the downstream subnetwork. More generally, this type of directional influence would tend to arise between any definable subnetworks with asymmetric interconnection strengths, i.e. where one subnetwork exerts a stronger influence than it receives in return. For visualization purposes, we demonstrated this concept using small-scale networks, though the underlying principles naturally extend to higher dimensional neuron populations. A concrete example of an upstream population controlling a downstream one is the communication between cortex to the spinal cord, which we will explore next.

### Emergent corticospinal control through (sub)network independence

In the brain, global network operation may need to be coordinated in such a way that many different subnetworks are controlled. A prime example is the spinal cord circuitry, which is under the control of the corticospinal system. The corticospinal system projects directly to the spinal interneuron population, which lacks a corresponding direct control of the corticospinal neuron population (the spinal interneurons instead predominantly send their ascending projections to the cerebellum (Jorntell 2017)). However, the spinal cord network inherently forms a vector field, both due to its intrinsic connectivity (Kohler et al 2022) and through its participation in sensorimotor loops via the periphery. By controlling its own activity state, the corticospinal system can alter the vector field that governs the dynamics of the spinal interneuron population. The movements initiated by that population generate sensory feedback which in turn can also influence the vector field impacting the spinal neuron population, because the sensory feedback will act as a third partially independent subnetwork, which has strong synaptic impacts on the spinal interneuron network. Therefore, in the presence of dynamic sensory feedback, which is always the case when there is actual body movement, dynamic movements can be produced even with limited impact from the corticospinal system.

But here, we were first interested in the state control exerted by the corticospinal system. Therefore, we connected 4 corticospinal neurons to a highly simplified spinal cord network (Figure 3A). Two neurons in the spinal cord constituted the readout population, i.e. these were two motorneurons that innervated different muscles. The corticospinal inputs consisted of tonic activity levels, with slight variations in the recruitment order across the input channels (Figure 3B). As shown in Figure 3C, a given input combination generated an attractor trajectory that the alpha-motorneurons tended to follow. The motorneurons produced a specific pattern of activation of the agonist and antagonist muscles, which was largely independent of the initial state. When the same corticospinal input pattern was played in reverse, this caused a different pattern of muscle activation, i.e. a different behavioral output, which was again robust to the initial state of the motorneurons (same set of initial states in Figure 3D,E). Hence, in the same given spinal interneuron network, without any intrinsic weight intrinsic changes, small variations in the corticospinal input can produce different spatiotemporal muscle activation patterns.

**Figure 3.**
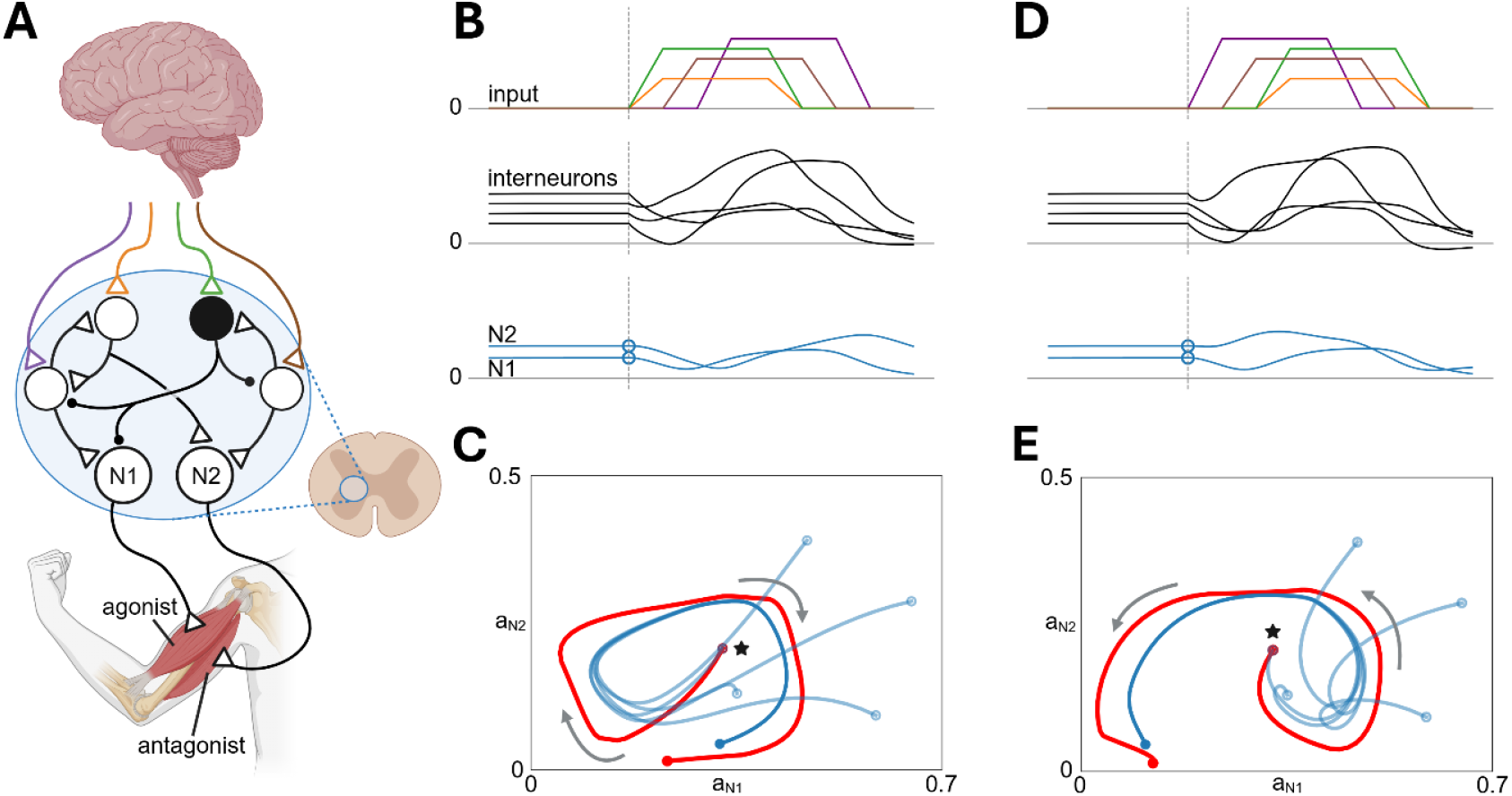
Corticospinal tract control of movement by biasing the state of the spinal cord network. A) The simplified spinal cord network, its control from a corticospinal input and its output to two muscles via alpha-motorneurons N1 and N2. B) The cortical inputs across four axons (color coded in A), the resulting activity across the spinal interneurons and motorneurons (N1, N2). C) The attractor trajectory and the activity trajectories of the motorneurons from different starting states. The attractor trajectory is in this case determined by the interneuron (upstream) subnetwork. Star symbol indicates the motorneuron activity trajectories shown in B. D,E) Same as in B,C but for a different cortical input.

This type of effect is not limited to the corticospinal impact on the spinal cord. For example, within the cortex, in any case where there is an unequal synaptic impact between two populations of intrinsically connected neurons, one can think of each population as a subnetwork where one subnetwork will tend to control the attractor trajectory in the other subnetwork. The more unequal the synaptic input between the two population, the more pronounced the subnetwork effect. This will create a “chasing its own tail” effect, where the state of the downstream population follows the attractor trajectory determined by the upstream population. This effect can be concatenated across an endless number of subnetworks and will determine how the cortex impacts its own state evolution, which can lead to what we think of as being equal to imagination, and anticipation. Notably, subnetworks do not need to be fixed entities, but their allocations and roles can instead change dynamically depending on state. Indeed, the functional connectivity, i.e. basically the correlation structure of neural activity, across the cortex rapidly changes in such a way that it is fully compatible with the dynamic subnetwork parcellation (Benisty et al 2024) that we describe here.

## Discussion

Here we used the vector field approach as a ‘ground truth’ framework to describe in principle how neuronal networks in the brain self-govern their time-evolving activity. We recently published the basics of the conceptual framework of vector fields to understand the interactions between neurons in globally interconnected networks (Szeier & Jörntell 2025), but in order to also explain the dynamics of network operation we here extended the vector field framework with the emerging effects following the incorporation of neuronal state memory. A dynamic state memory is present in each neuron due to the presence of the neuronal membrane capacitance and the resulting dynamic leak. We find that a continually evolving global neuron activity distribution as observed in the brain *in vivo* arises when the global network is subdivided into subnetworks, where a subnetwork is defined as a group of interconnected neurons with unequal impacts on another subnetwork. We illustrated this principle with the corticospinal control of the spinal cord but underscore that subnetworks will also dynamically arise across the cortex, as many observations imply (Benisty et al 2024, Etemadi et al 2022, Etemadi et al 2023, Norrlid et al 2021). In both cases, the emerging effect is that an upstream subnetwork will dynamically move the apparent attractor point in the downstream subnetwork, thus creating an attractor trajectory in the downstream subnetwork, which will exert a powerful control over the temporal evolution of its neuron population activity.

### Importance to explain behavioral control

The corticospinal control of the spinal cord is the ultimate step in the regulation of purposeful, non-reflexive behavior. We showed how a relatively simple corticospinal signals can result in complex attractor trajectories in the spinal cord network. Because the activity of the spinal cord tends to follow whatever trajectory the attractor follows, this can in turn result in diversified movement patterns. Small adjustments to the corticospinal signal structure, which should be relatively easily learnt by the cortex even from early development, can lead to quite different movement patterns. Hence, an advantage of this approach is that it can facilitate gradual trial and error motor learning in the cortex. Small differences in corticospinal output can lead to large differences in motor output, due to the attractor-based functionality in the target network, which will tend to amplify smaller differences. Hence, the attractor trajectories that arise due to this dynamic vector field control can explain how a diversity of motor behaviors arises, even though the same network is engaged across all the movements in the examples given.

Although not shown directly, the sensor feedback to the spinal cord (Kohler et al 2022) can be viewed as another subnetwork input to the spinal cord. Sensory feedback will therefore also impact the attractor trajectory in the spinal cord. This will stabilize the current dynamic solution in the spinal cord circuitry, making it more of a self-playing piano at least until the descending excitation of the spinal cord stops. Other factors that would have an impact on the spinal cord circuitry activity are the brainstem nuclei with descending influences. The brainstem nuclei are more reactive systems, i.e. the vestibular sensors will directly drive the vestibulospinal nuclei which will impact the spinal cord activity. By extension of the same principle, this would influence the attractor trajectory in the spinal cord circuitry in much the same way as the sensory feedback. Then we can think of the cerebellum as a read-out of the state in the spinal interneuron population and forward adjusting the activity in the spinal cord in accordance with whatever principles is assumed to govern the internal connectivity of the cerebellum (Jorntell 2017).

### Extension to more indirect behavioral control

Behavioral control is not only motor execution but also deciding when and if a particular movement pattern is to be executed. The cortex is an important factor in this more indirect process, not the least because its ongoing activity is in part reflective of a working memory (Kristensen et al 2024). In this case, the cortex must regulate its own activity to determine the appropriate timing and content of state bias signals sent to the spinal cord, thereby increasing the likelihood of generating a specific movement pattern. We demonstrate, in principle, how this cortical self-governing can operate. Naturally, in the real brain, the structure of the network is gradually adapted by sensory experiences (Ebrahimi et al 2022), which helps refine cortical self-governing to better support real-world interactions. The mechanisms underpinning self-governing are illustrated here. The thalamus and the basal ganglia exert their influences on the cortex, hence they will bias the attractor trajectories across the cortical subnetworks.

### Implications for anticipation and illusion

Our framework can explain anticipation. Anticipation can be demonstrated by that the exact same sensory input can yield widely different responses in the cortical network (Etemadi et al 2022, Norrlid et al 2021) and that isolated episodes of brain activity at rest often resembles that during behavior or sensory input patterns (Jafari et al 2020, Luczak et al 2009, Norrlid et al 2021). Anticipation can only originate from network learning from previous monitoring and trying to understand the sensory world. As the anticipation becomes better and more reliable, we can gradually scale down the number of sensory cues that we need to predict how the world is going to evolve. Eventually, when we can do that without any cues at all it becomes imagination of future sensory input (Jafari et al 2020).

This framework can straight-forwardly explain cross-modal illusions or other types of illusions. I.e. consider if we are currently on an attractor trajectory for executing a certain movement and then there is visual information to the visual cortex. The visual information is in this case the upstream subnetwork that can impact the activity of the downstream subnetwork, with more corticospinal neurons, such that its attractor trajectory changes path, and therefore the movement executed will be impacted. The cortical system shares information across all its subnetworks (Ebrahimi et al 2022, Enander et al 2019, Etemadi et al 2022, Etemadi et al 2023, Khilkevich et al 2024, Mimica et al 2023, Nietz et al 2022, Wahlbom et al 2019, Wahlbom et al 2021), so we can naturally replace visual information in the example by any other modality or internal representation that is being engaged. We can think of perception as being an attractor trajectory with similar properties, except that it doesn’t necessarily result in overt movement. Rather, it serves as an internal recording of an event, which can later influence our behavioral choices. In the case of an illusion, this recording is shaped by visual information in a misleading way. In general, our framework can explain illusions in the following way: an anticipatory attractor trajectory, established prior to the arrival of sensory information, can be influenced by contextual cues. These cues stem from representations within the upstream network, which impacts the vector field of the downstream network (the ‘readout’ population).

### Limitations

We introduced this framework of vector field dynamics to better understand the emergent operation in neuronal networks, how the activity distribution across its neuron population will gradually alter solely depending on intrinsic factors. Are there situations when this framework would become non-applicable, or less relevant? In the presence of recursive connectivity, the network begins to exhibit potentially complex dynamics, where the function of a given neuron can potentially alter dramatically given the state of the other neurons of the network. This is the situation where we believe this framework is the most useful. In contrast, in pure feed-forward networks, where the neurons are organized in layers and there are no interconnections within a layer, while the dynamic vector field framework is still applicable to describe such a network it may not add substantial understanding compared to other methods to describe the properties of such networks. However, within the brain, recursive connectivity exists at least within the cortex, the thalamocortical system and within the spinal cord circuitry, as well as in the spinal cord sensorimotor loops through the body. Recursive connectivity may be less prominent in brainstem nuclei and the cerebellum, but these structures exert their functions via the former systems. Under these circumstances, the output function of any given neuron within the central nervous system will change dynamically and this is an emergent effect that is currently poorly accounted for in neuroscience theory and where the present framework could be helpful to analyze and understand such phenomena.

### Robust operational principle that can support evolutionary development

The examples presented were obtained under specific parameter settings in the neurons, with their time-dependent activation function, and specific connectivity between the neurons. The sensitivity to adjustments in these parameter settings can be expected to be low, as we previously showed that different activation functions and thresholds will still produce the overall vector field structures that we used here to simulate the dynamic behavior of small circuits (Szeier & Jörntell 2025). This is one of the powers of the approach, that its overall functionality is robust to large variations in neuron behavior, number of synaptic inputs and synaptic weights. The regulation of the synaptic weights, highly skewed or more uniform, can be expected to greatly impact the relative contribution of individual neurons to the dynamics of the network behavior, but will not change the operational principle. Being an emergent effect from how individual neurons work and are interconnected, the principles of this foundational conceptual framework could hence apply to explain the basis of any behavior across any species that depend on a nervous system, and the evolutionary addition of additional CNS structures would merely feed into this conceptual framework as subnetworks biasing the state of the rest of the network.

## Acknowledgements

This work was supported by VINNOVA and the Swedish Research Council (VR).

